# Engineering an in vitro model of demyelinated spinal cord tissue

**DOI:** 10.64898/2026.02.15.706037

**Authors:** Lu Jin, Natasha Brinkley, Youyi Tai, Galilea Flores, Jin Nam

## Abstract

Demyelinating diseases are a group of complex neurodegenerative disorders characterized by damage to the myelin, the protective sheath that insulates and supports efficient nerve signal conduction. Such a loss of myelin causes the formation of lesions not only in the brain but also often in the spinal cord (SC). Despite the high prevalence of SC lesions among patients, existing models mostly focus on those in the brain, inadequately capture the unique anatomical and physiological features of SC pathology. In this study, we developed a robust, reproducible in vitro model of SC demyelination by combining microwell technology and piezoelectric scaffolds to engineer human neural stem cell (hNSC)-derived nerve tissues featuring aligned, myelinated, extended axons up to 2000 µm in length. We utilized distinct chemical treatments to induce demyelination with or without axonal degeneration: a cuprizone cocktail, a copper chelator combined with inflammatory cytokines, and lysophosphatidylcholine (LPC). Electrophysiological assessments validated the physiological relevance of our model, demonstrating impaired signal transmission and neural connectivity akin to in vivo demyelination pathology. Our versatile platform thus provides a valuable tool for elucidating SC demyelination pathophysiology and exploring potential therapeutic interventions.

## 1. Introduction

Demyelinating diseases are a group of neurological disorders that the myelin sheaths surrounding axons are damaged, leading to disruptions in nerve signal transmission. By insulating the axon and preventing ion leakage, the myelin sheath facilitates saltatory conduction, a process where the nerve impulse rapidly jumps between unmyelinated gaps (nodes of Ranvier) rather than moving continuously along the axon [1]. Thus, demyelination compromises axon transmission functionality [2], and potentially results in axon degeneration [3]. These demyelinating diseases include multiple sclerosis (MS), neuromyelitis optica spectrum disorder (NMOSD), acute disseminated encephalomyelitis (ADEM), and Guillain-Barré syndrome (GBS), affecting both the central nervous system (CNS) and the peripheral nervous system (PNS).

Demyelinating diseases are multifactorial disorders influenced by genetic predisposition, environmental exposures, and lifestyle, which collectively contribute to immune system dysregulation. MS is the most common demyelinating disease in the CNS, affecting approximately 2.8 million people worldwide. It is characterized by demyelination, oligodendrocyte loss, multifocal inflammation, axonal degeneration, breakdown of the blood-brain barrier, and reactive gliosis [4, 5]. Among them, demyelination lesions are the primary characteristic of MS pathological hallmark, and they are involved in both the white mater (WM) and grey matter (GM) [6, 7]. Approximately 80-90% of MS patients develop SC lesions in both brain and spinal cord (SC) regions [8, 9], leading to symptoms such as impaired movement, coordination issues, and dysfunction in bladder and bowel control [5, 10, 11].

Despite this high incidence of the involvement of the SC in demyelinating diseases, studies on SC-specific pathology remain limited compared to those focusing on brain pathology, particularly regarding anatomy, pathophysiology, the mechanisms of disease progression, and potential disease-modifying treatments (DMTs) [7]. Given the complexity of SC pathology, significantly different from that in the brain, further studies are necessary to better characterize and understand the mechanistic details of demyelinating processes and their pathological outcomes in the SC. The main challenge in studying demyelination in the SC, however, is the difficulty of imaging SC regions due to their small sizes, anatomical variability in different interfaces, and motion artifacts [7, 11–15].

Currently, a variety of models, including animal proxies, ex vivo, and in vitro platforms, are employed to mimic the pathogenesis of demyelinating diseases. While most insights on the primary mechanisms of disease progression have been derived from preclinical animal studies and their organotypic cultures [16–18], these models fail to fully replicate human pathophysiology, limiting their translational value [19]. In this regard, in-vitro models based on human cells could recapitulate the physiopathology specific to humans. With the limitation of maintaining cell viability and identity under co-culture conditions of primary cells, human induced pluripotent stem cells (iPSCs) or neural stem cells (NSCs) provide a viable platform for co-culture of multiple cell types derived from a single origin. It is worthwhile to note the recent progress in developing spheroids and organoids to further elucidate the intricate cell interactions [20–22]. However, these in-vitro models predominantly focus on brain-specific aspects of demyelinating diseases instead of spinal cords, partly due to the challenge of replicating physiological relevance, including extended uniaxial axons, cellular diversity, and adequate myelination.

In our recent studies, we established the potency of mechano-electrical stimulation (MES), mediated by piezoelectric electrospun poly(vinylidene-trifluoroethylene) ((P(VDF-TrFE)) nanofibrous scaffolds, to induce the multiphenotypic differentiation of hNSCs simultaneously toward neurons and glial cells, resulting in the formation of mature neural tissues [23, 24]. These results provide the potential to develop various in vitro neurological disease models by leveraging the development of multiple, coexisting phenotypes from the single origin of hNSCs. Specifically, this approach enables intricate interactions among various neural cell types, crucial for disease pathogenesis and progression in demyelinating diseases. In order to offer more physiologically relevant insights into demyelination in SCs, control over cellular organization such as uniaxial alignment of long axons is critical as demyelination in such a structure could amplify the signal transmission defects over a long distance unlike the brain [25]. However, materializing physiologically relevant extended, myelinated axons in vitro remains challenging. To address this, we developed a microwell system to inoculate cells in a spatially controlled manner, combined with a temporally varied biochemical supplementation, to promote directional axonal growth and elongation with mature myelination. We then explored and compared the pathology in engineered nerves under chemically induced demyelination by various biochemicals, including cuprizone cocktail and lysophosphatidylcholine (LPC, also known as lysolecithin), by morphological and electrophysiological analyses. This comprehensive approach established a highly reproducible in vitro model that mimics the morphology of axons in the SC, with electrophysiological relevance. We expect this in vitro model to advance our understanding of demyelination in SCs and ultimately guide the development of improved diagnostic and therapeutic strategies.

## 2. Materials and methods

### Scaffold synthesis and characterization

Electrospun P(VDF-TrFE) nanofibrous scaffolds were synthesized as previously reported [23]. Briefly, 17.0 wt% polymer solution was prepared by dissolving 70:30 mol% P(VDF-TrFE) copolymer (PolyK) in a solvent mixture consisting of 60/40 volume ratio of dimethylformamide (DMF, Sigma Aldrich) and acetone (Thermo-Fisher), and 8.0 wt% pyridinium formate buffer (PFB, Sigma Aldrich). The solution was electrospun in an enclosed system featuring a grounded rotating collector drum for aligned fiber deposition. Optimized electrospinning parameters included a tip-to-collector distance of 10 cm, a voltage application of –15 to – 20 kV at the tip and + 0.1 kV at the collector, a flow rate of 6 mL/h, a collector rotation speed of approximately 47.9 m/s at ambient conditions of 23 °C with an absolute humidity of 10.283 g/m³ (∼50% RH). The electrospun fibers were collected for 3 hrs. These optimized parameters yielded a scaffold with an average fiber diameter of 500 nm and a thickness of approximately 200 µm. Afterwards, the scaffolds underwent annealing at 90 °C for 12 hrs to enhance crystalline phase transition, as a result, further improving piezoelectricity [26] and being ideal for inducing hNSCs differentiation under MES [23]. Scaffold morphology and fiber diameter were characterized via scanning electron microscopy (SEM, VEGA3, Tescan, Czech Republic) with the ImageJ software (NIH).

### Microwell mold fabrication

Microwell molds utilized for spatially controlled cell inoculation onto the electrospun scaffolds were fabricated from polydimethylsiloxane (PDMS) through a standard mold-casting technique. The microwell mold was designed using the SolidWorks software (Dassault Systèmes, France), and 3D-printed using a Digital Light Processing (DLP) Craftsman resin (Anycubic, Hongkong) with an Anycubic Photon Ultra stereolithography (SLA) resin printer. SLA printing was employed over filament extrusion due to its superior capability in achieving the micron-scale precision required for our microwell design. Furthermore, to prevent PDMS adhesion to the resin mold, a permanent non-stick surface coating was applied with a thin layer of photoresist (SU-8, Microchem) on the surface of the resin mold. The degassed PDMS solution was poured onto the prepared resin mold, spin-coated at 1200 rpm for 20 secs to achieve uniform mold thickness, and cured at 60 °C for 1 h. Once cured, the PDMS microwell mold was carefully peeled off from the resin mold and subjected to a three-step cleaning process, including sonication for 10 mins sequentially in acetone (Thermo-Fisher), 70% ethanol (Decon), and deionized water to remove surface contaminants. Following cleaning, the microwell sheets were manually sectioned to match scaffold dimensions, treated with atmospheric plasma (Harrick Plasma) to enhance hydrophilicity, and sterilized by autoclaving prior to use.

### Mechano-electrical stimulation-induced neuromorphogenesis of human neural stem cells under temporally controlled biochemical supplementation

All experiments involving hNSCs were approved by UC Riverside Institutional Review Board (IRB; HS11–124) and Stem Cell Research Oversight (SCRO; SC20210002) Committee. A well-characterized hiPSC line was utilized to derive hNSCs as described elsewhere [23]. NSCs were cultured on Geltrex (Gibco)-coated tissue-culture plates (Thermo-Fisher) in neural maintenance media (49.5% Neural Basal Media + 49.5% Advanced DMEM F-12 Media + 1% Neural Induction Supplement, Gibco) at 37 °C under 5% CO₂. P(VDF-TrFE) scaffolds were cut into 5 × 40 mm strips, with both ends coated with hydrophobic poly(styrene-block-isobutylene-block-styrene) (SIBS, Sigma-Aldrich) elastomer to define a 5 × 10 mm cell culture area. The scaffolds were sterilized with 70% ethanol, rinsed with phosphate-buffered saline (PBS, Corning), surface treated with poly-L-ornithine (Sigma-Aldrich) diluted in borate buffer (Thermo-Fisher) for 1 h, and then finally coated with laminin (Thermo-Fisher) diluted in PBS for 2 hrs to enhance cell adhesion and differentiation. After the protein coating, the PDMS microwell mold was attached to the scaffold assembled in a cell culture chamber and allowed to anneal overnight [23]. On the next day, hNSCs were seeded into the microwells at a density of approximately 100,000 cells/cm² scaffold area. After allowing the cells to settle and adhere to the scaffolds for 30 mins, the mold was removed and neural maintenance media were added into the culture chamber. Cells were subject to MES, as described elsewhere [23, 24], for the first four weeks to facilitate neural differentiation and maturation. In addition, after cell seeding on Day 0, 50 ng/mL NGF (Sigma-Aldrich) and 10 µM ROCK inhibitor (Y-27632, STEMCELL Technologies) were supplemented into the cell maintenance media from Day 1–4, and the concentrations were further increased to 100 ng/mL and 20 µM, respectively from Day 5–14 to enhance axon elongation while inhibiting glial cell differentiation [27]. Afterwards, cells were subjected to culture for 2 additional weeks under MES in the maintenance media to recover glial cell differentiation and myelination.

### Biochemically induced demyelination of engineered nerves

The engineered nerve tissues cultured for 4 weeks under MES and sequential biochemical treatment as described above, were then treated with demyelination reagents of either 1) a cuprizone cocktail composed of cuprizone (Sigma-Aldrich) at 1000 µM combined with inflammatory cytokines including 50 ng/mL TNF-α and 50 ng/mL IFN-γ (both from Proteintech) or 2) LPC (Sigma-Aldrich) at 30 µg/ml for 24 hrs before switching back to the normal maintenance media. Engineered tissues were harvested at different time points as necessary for immunostaining or electrophysiology analysis.

### Immunofluorescence imaging

Samples were fixed using 4% paraformaldehyde (PFA, Sigma-Aldrich) for 30 mins, rinsed with PBS at 4 °C. For immunofluorescence (IF) staining, the fixed samples were permeabilized with 0.1% Triton X-100 (Thermo-Fisher) for 30 mins, and blocked with 1% bovine serum albumin (Sigma-Aldrich). Primary antibodies were added to the samples, following vendor-recommended dilutions in 0.02% Tween 20 (Sigma-Aldrich) in PBS (i.e., PBST) and incubated overnight at 4°C. Primary antibodies used in this study include Tuj.1, a β3-tubulin antibody (Proteintech) for axon/neuron; Galactocerebroside (GalC, Proteintech), myelin oligodendrocyte glycoprotein (MOG, Santa Cruz Technology) and myelin basic protein (MBP, Bio Rad) for oligodendrocyte and myelin; excitatory amino acid transporter 2 (EAAT2, Santa Cruz Technology) and glial fibrillary acidic protein (GFAP, Cell signaling) antibodies for astrocytes; vesicular glutamate transporter 1 (VGLUT1, Thermo-Fisher) and postsynaptic density protein 95 (PSD95, Thermo-Fisher) for excitatory synapse; and vesicular gamma-aminobutyric acid (GABA) transporter (VGAT, Thermo-Fisher) and gephyrin (Thermo-Fisher) for inhibitory synapse. After overnight incubation, samples went through a thorough washing with PBST, and stained with appropriate secondary fluorescence antibodies (Abcam) for 1 h, rinsed with PBST, followed by mounting with Vectashield containing DAPI (4’,6-diamidino-2-phenylindole, Vector Labs). The stained samples were observed under a Zeiss-880 upright confocal microscope. The ImageJ software was utilized to quantify cell number, axonal length and anisotropy, and the degree of demyelination.

### Electrophysiological assessment of demyelinated neural networks

The effects of demyelination on the neural signal transmission were assessed by measuring the electrophysiological activities of engineered neural networks using a multielectrode array (MEA; MultiChannel Systems, Germany). The MEA comprised 60 titanium nitride (TiN) electrodes (30 µm diameter) arranged in an 8 × 8 grid with 200 µm spacing, excluding the four corner electrodes. After neural tissue formation and subsequent demyelination treatment, the engineered tissues were carefully removed from the culture chambers and placed upside-down onto the MEA surface, facing cells to the electrodes and ensuring orthogonal alignment of axons across the patterned electrodes. The extended axons across the two colonies formed by the microwells enabled neural signal transmission from one side of MEA electrodes to the other side. A small weight of approximately 10 g was applied to maintain consistent and direct electrode-cell contact. To evaluate neuronal connectivity and network function, one end row of electrodes was selected to deliver a stimulation with a biphasic pulse of +/−500 mV amplitude, 200 µs duration per phase, repeated 10 times with 2 sec intervals. Evoked responses (i.e., action potentials) were recorded from adjacent electrodes, enabling analysis of network connectivity, signal transmission distance, velocity along the long axons between the two cell colonies. During measurements, the sample was maintained in the neural maintenance media within an incubator at 37 °C in 5% CO₂.

### Statistical Analysis

All experiments were conducted with at least biologically independent triplicates. Statistical analysis was performed using one-way ANOVA followed by a post-hoc test (Tukey’s test) with the SPSS software (IBM). Significant differences are indicated by asterisks, i.e., * for *p* < 0.05 and ** for *p* < 0.01. Results are presented as the mean ± standard deviation. All quantification plots were generated using the Origin software (OriginLab).

## 3. Results and Discussion

### Microwell Fabrication

Previous work from our lab successfully demonstrated the differentiation of hNSCs simultaneously into three major neural cell types (neurons, astrocytes, oligodendrocytes) by the application of MES using piezoelectric electrospun scaffolds [23, 24]. However, deriving a physiologically relevant length of extended axons to simulate the SC remains challenging. To guide such structural formation, we incorporated a microwell system for precise cell inoculation to form spatially controlled cell colonies and to enable the extension and alignment of axons between those colonies. A microwell cell inoculation mold was fabricated and refined through multiple iterations (Fig. S1). A 3D printed positive resin mold having the microwell dimensions of 1200 µm × 400 µm × 500 µm (L × W × H), separated by a gap distance of 500, 1000, 1500, or 2000 µm was designed, coated with photoresist (Fig. 1A-B). PDMS was casted in the mold by spin-coating (Fig. 1C). After curing, the PDMS microwell mold was peeled off gently (Fig. 1D), and cut into 0.5 × 4 cm strips (Fig. 1E). Following plasma treatment (Fig. 1F) and sterilization, the microwell molds were placed face-down onto the scaffolds (Fig. 1G), and assembled into custom culture chambers (Fig. 1H). The plasma treatment ensured the mold remained adhered to the scaffold for cell seeding, yet they could be still removed from the scaffold later without damaging the cells/scaffold. Maintaining a tight seal between the molds and scaffolds was critical for achieving the formation of uniform cell colonies (Fig. S2); plasma treatment on the inverted mold surface (Inv/PT) exhibited the optimized cell colony shape consistency, confirmed by IF imaging, whereas other conditions resulted in non-uniform, disorganized cell distribution. After cell attachment for 30 mins, the molds were removed from the scaffolds and the cell culture chambers placed within a 6 well-plate were positioned on a vertical translation stage as described elsewhere [23, 24]. The periodic vertical motion of the stage delivered mechanical stimuli to the piezoelectric scaffolds, and the scaffolds exerted concurrent electrical stimuli to the cells via the piezoelectric effect to realize the MES effect. The scaffold deflection is optimized to elicit a piezoelectric response measured at approximately 200 mVp-p, ideal for hNSC differentiation and mature tissue formation [23].

**Figure 1.**
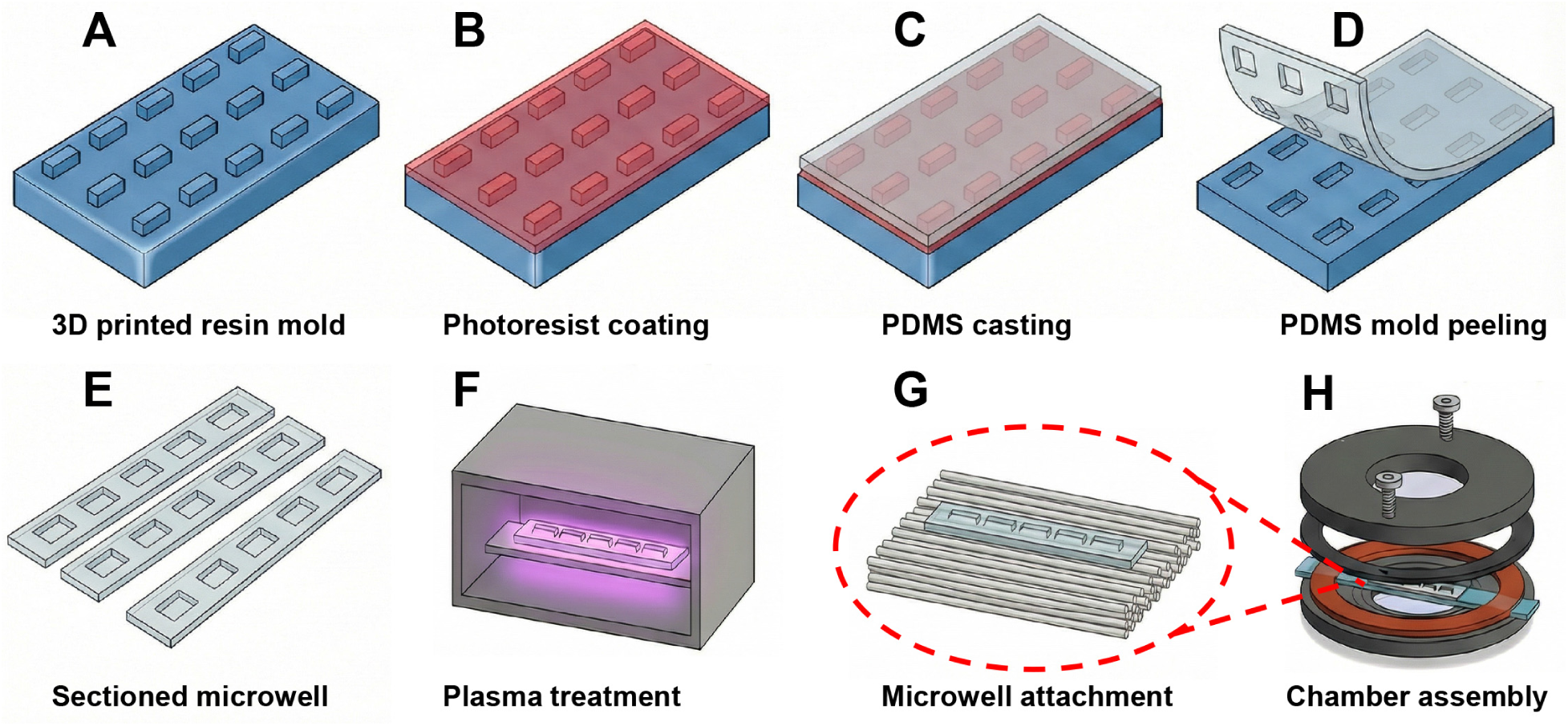
Design and fabrication of PDMS microwell molds. (A) Optimized positive resin mold fabricated by 3D printing. (B) Photoresist on the surface of the resin mold. (C) PDMS casted on the 3D-printed resin mold by spin coating. (D) PDMS microwell mold peeled from the resin mold. (E) Sectioned microwell mold for controlled cell inoculation. (F) The scaffold adhesion side of the microwell mold being subjected to plasma treatment. (G) Microwell mold placement on electrospun scaffold. (H) Cell culture chamber assembly with the microwell mold-attached scaffold.

### Uniaxial and long axon extension with myelination

To enhance axonal extension to mimic the highly polarized architecture of the SC [28, 29], the culture medium was supplemented with NGF and ROCK inhibitor for the first two weeks under MES [27]; these biochemical factors were incorporated to promote axonal growth while preventing the differentiation of hNSCs to myelinating glia, which have been shown to significantly inhibit neurite outgrowth [30, 31]. MES was initiated two days post-seeding and maintained for the duration of the culture period. As depicted in Fig. 2A-D, distinct cell colony formation and robust axonal bridging between such colonies were induced by microwell seeding and subsequent culture under MES for all gap distances of 500, 1000, 1500, and 2000 µm. After achieving the axonal bridging within the initial 2-week culture duration, NGF and ROCK inhibitor were withdrawn from the culture medium to allow glial differentiation and maturation over the following weeks. Since Week 3, the oligodendrocyte differentiation of hNSCs and progressive myelination were observed within the colonies and gaps across the colonies, as shown by triple immunostaining of β3-tubulin, an axonal marker, GALC and MBP, intermediate/mature markers for myelination, respectively (Fig. 2E-J, K-P). Both regions within and across colonies showed vigorous neural network formation while GALC was expressed as early as Week 3 indicating progressive oligodendrocytic differentiation. In addition, oligodendrocytic maturation and myelin sheath formation were further supported by a time-dependent increase in GALC/β3-tubulin and MBP/β3-tubulin ratio (Fig. 2Q-T).

**Figure 2.**
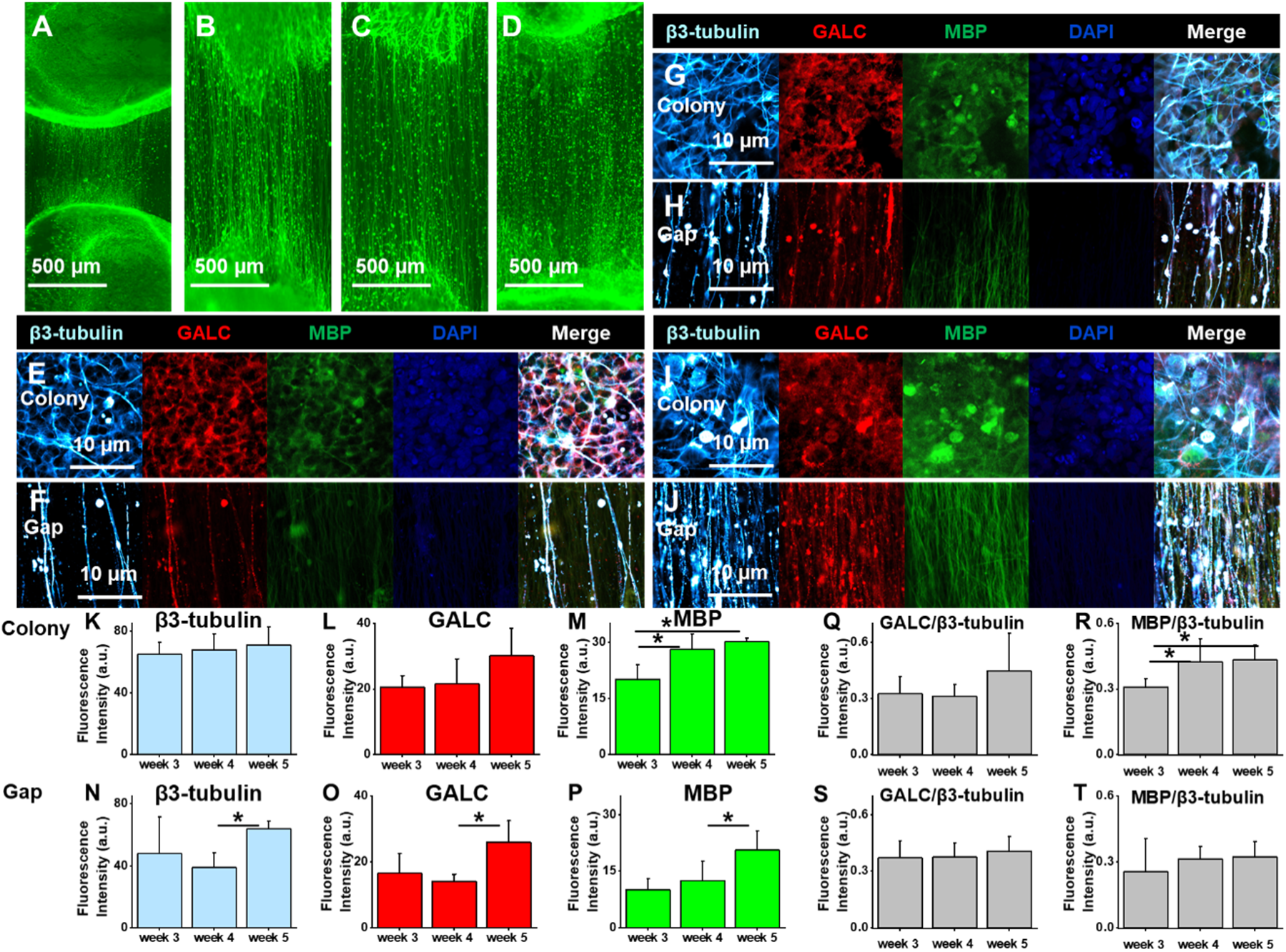
Optimized extension of axons bridging cell colonies and their myelination. (A-D) Development of aligned axons (labeled by β3-tubulin) bridging cell colonies formed by microwell mold cell seeding with various gap distances of 500 µm (A), 1000 µm (B), 1500 µm (C), and 2000 µm (D). (E–J) Development of neural network (β3-tubulin) and myelination (early stage myelination by GALC and late stage by MBP) within the cell colonies (E, G, I) and inter-colony gaps (F, H, J) cultured for 3 (E, F), 4 (G, H) and 5 weeks (I, J) under mechano-electrical stimulation. (K-T) Quantification of axonal development and myelination in (E-J) based on marker intensities including β3-tubulin (K, N), GALC (L, O), MBP (M, P), and myelination per axon by ratio of GALC/β3-tubulin (Q, S), MBP/β3-tubulin (R, T) within the cell colonies (K-M, Q, R) and inter-colony gaps (N-P, S, T) over 3 weeks (n=3, \**p* < 0.05, \*\**p* < 0.01).

Since robust axonal bridging was achieved in the largest gap distance tested at 2000 µm, this culture condition was utilized in the subsequent experiments. A full-scale overview of axonal extension with mature myelin formation across adjacent colonies are shown in Fig. 3A. The axon outgrowth was highly unidirectional, with a mean orientation angle of 90.21°, and the distribution of orientation angles closely approximated a normal distribution (Gaussian R² = 0.98) (Fig. 3B). In addition to achieving long and unidirectional axonal alignment, ensuring their maturity and functionality is also critical for an in vitro model to accurately simulate in vivo nerve tissues with physiological relevancy. Therefore, the functional neural network formation in the engineered tissues was determined by examining excitatory and inhibitory synapses (Fig. 3C-D). Excitatory synapses were labeled by pre-synaptic VGLUT1 and post-synaptic PSD95, with their colocalization indicating active puncta formation. Both the colony region and the gap region exhibited such puncta formation, indicating the presence of synapses within the colonies and along the extended axons. A zoomed-in picture from long extended axons further signified the excitatory puncta formation, potentially indicating active signaling between the colonies. Inhibitory synapses were similarly identified by VGAT and Gephyrin expression (Fig. 3D). The existence of both excitatory and inhibitory synapses confirmed the formation of a complex, functionally mature neuronal network. Moreover, some regions displayed axoplasmic fusion and thick neurite bundle formation, potentially suggesting SC–like organization, supported by strong MOG expression marking the presence of mature oligodendrocytes (Fig. 3E). Thus, by implementing MES with the sequential biochemical treatment of NGF and ROCK inhibitor on hNSC colonies formed by microwell-guided cell inoculation, we successfully engineered nerve tissues characterized by extended, unidirectional axon outgrowth over a 2000 µm gap with extensive myelination and functional synapse formation from a single origin of hNSCs.

**Figure 3.**
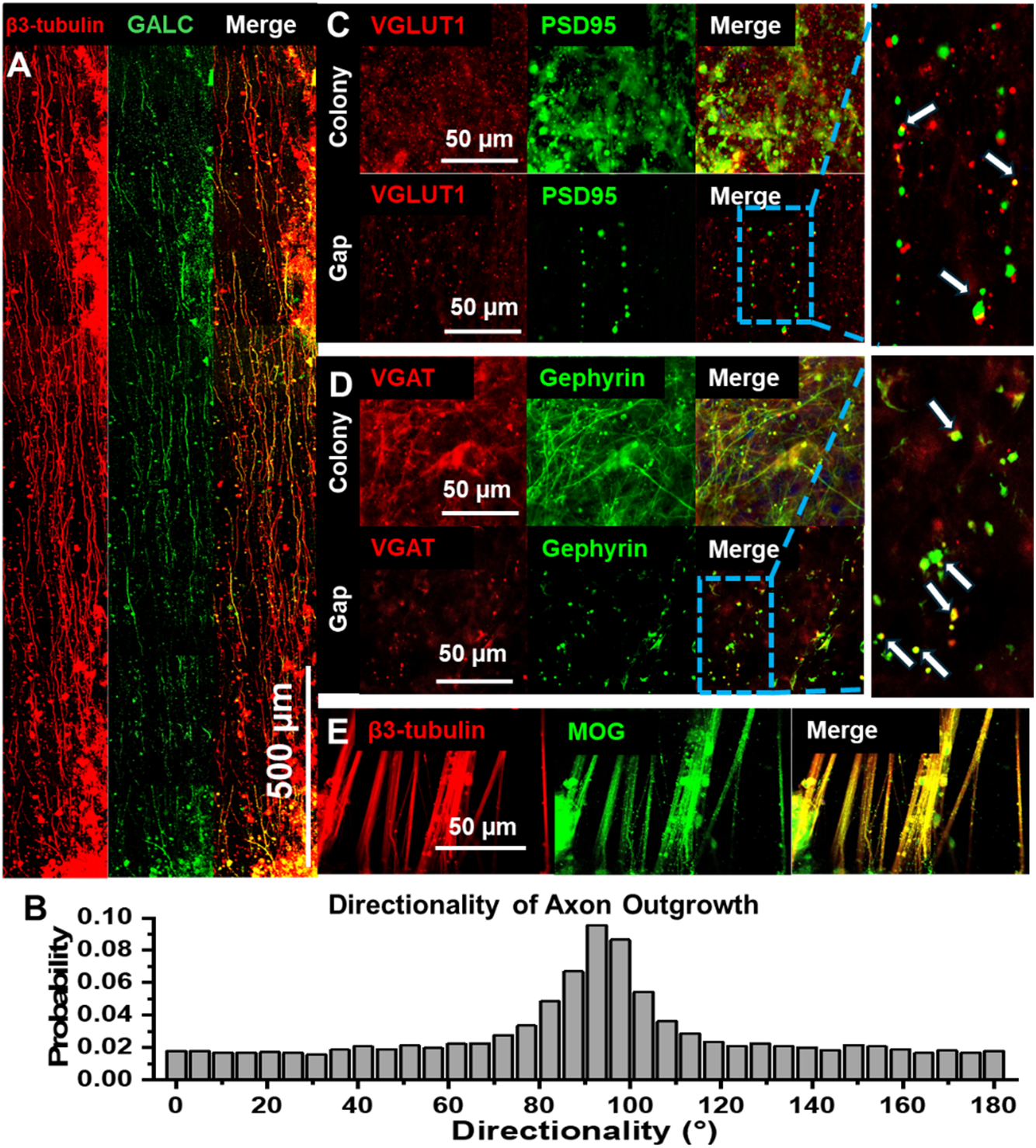
Development of uniaxially aligned, myelinated axons with functional synapse formation. (A) Formation of myelinated axon extension between colonies formed by a microwell mold having a gap distance of 2000 µm. (B) Histogram of axon directionality. (C-D) Formation of the excitatory (C) and inhibitory synapses (D) determined by pre- and post-synaptic markers, including VGLUT1/PSD95 (excitatory) and VGAT/Gephyrin (inhibitory), within the colonies and inter-colony gaps. (E) Formation of nerve bundles among the extended mature axons, indicated by the groups of nerve fibers (β3-tubulin) with MOG expression for mature myelination.

### Demyelination of the engineered nerve tissues

Leveraging this engineered myelinated nerve tissue composed of neurons and glial cells, mimicking the structure of the SC, we next investigated its potential as a model for demyelinating diseases. To induce demyelination, we tested a cuprizone cocktail (cuprizone + inflammatory cytokines, including TNF-α and IFN-γ) and LPC, both recognized for their demyelinating effects [32, 33]. For experimental demyelination in animals, cuprizone is the gold standard for toxicity-induced demyelination by selectively impairing oligodendrocytes and inducing oligodendrocyte apoptosis, although its precise mechanism remains unclear [34, 35]. Furthermore, the myelin sheath loss caused by cuprizone could diminish axonal integrity, resulting in permanent axon damage and loss [36, 37]. Interestingly, certain CNS regions exhibit differential susceptibility to cuprizone, with the SC tissue being notably resistant. Such findings underline existing limitations and emphasize the need for improved in vitro models specifically targeting SC demyelination. Due to the higher permeability of the blood-spinal cord barrier (BSCB) compared to the blood-brain barrier (BBB), inflammatory cytokines such as tumor necrosis factor-alpha (TNF-α, or Cachectin), interferon-gamma (IFN-γ), and interleukines (IL-1β, IL-6), are easier to diffuse into the cord regions, contributing to the progression of demyelination and inflammation [38]. Pasquini et al. reported negligible effects of cuprizone alone, noting significant oligodendrocyte disruption only when combined with pro-inflammatory cytokines TNF-α and IFN-γ [39]. TNF-α exerts pro-inflammatory effects through tumor necrosis factor receptor 1 (TNFR1) by damaging oligodendrocytes and the myelin sheath, subsequently leading to neuronal injury and axonal degeneration [40, 41]. In addition, Timothy et al. showed that IFN-γ induces oligodendrocyte cell death by apoptosis [42]. Another approach involves the direct administration of LPC into CNS tissue. LPC, an endogenous lysophospholipid elevated in the cerebrospinal fluid of MS patients [43], induces demyelination by increasing cellular membranes’ permeability, disrupting lipid membranes in the myelin sheath [44].

To induce demyelination while minimizing cell apoptosis, the optimal concentrations of these reagents were first determined by a cytotoxicity test (Fig. S3). Cuprizone alone at the maximum concentration of 3000 µM didn’t significantly decrease cell viability. At this concentration, however, cuprizone did not induce demyelination in this in vitro culture condition (Fig. S4). In contrast, exposure to 3000 µM cuprizone with TNF-α (50 ng/mL) and IFN-γ (50 ng/mL) drastically reduced cell viability. To maintain the cell viability, especially neurons, while maximizing the demyelination effect on myelin sheath, 1000 µM cuprizone supplemented by TNF-α and IFN-γ (Cup Cocktail) was selected for the demyelination treatment regime. The need of the inflammatory cytokine supplement for cuprizone-induced demyelination in vitro could be extrapolated from the fact that the effect of cuprizone in vivo is mediated by the inflammatory cytokines secreted by microglia/astrocytes/macrophages, resulting in demyelination [39, 46]. Similarly, 30 µg/mL LPC was selected as the highest effective concentration that preserved cell viability while inducing demyelination as any dosage beyond that severely caused cell apoptosis/necrosis (Fig. S3B, E). These reagents were introduced during culture in Week 4 for a short duration of 24 hrs, followed by a one-week recovery to stabilize demyelination (Fig. 4A). The samples were harvested for IF staining at Week 5. As expected, control samples without the application of demyelination reagents retained extensive axonal network with robust myelination based on strong MOG expression (Fig. 4B-C). Conversely, Cup Cocktail-treated samples exhibited significantly diminished MOG expression as well as disrupted neuronal network based on disconnected β3-tubulin distribution in both within cell colonies and gaps (Fig. 4D-E). These results indicated that Cup Cocktail induces axon degeneration as well as myelin loss. In LPC-treated samples, however, myelin loss was mostly localized to the gap regions while neuronal network remained relatively unaffected (Fig. 4F-G). Quantitative analysis further confirmed that both demyelination reagents damaged myelin sheath while LPC selectively induced demyelination in the uniaxially extended axons, based on the decreased MOG/β3-tubulin ratio in the gap regions as compared to that within colonies (Fig. 4H-N). In contrast, Cup Cocktail group degenerated neural network in both colonies and gaps. These results suggest that our engineered nerve tissue system can serve as a robust in vitro platform for demyelinating diseases by applying appropriate chemical treatments, capturing myelin loss and/or neuronal network disruption in a physiologically relevant condition.

**Figure 4.**
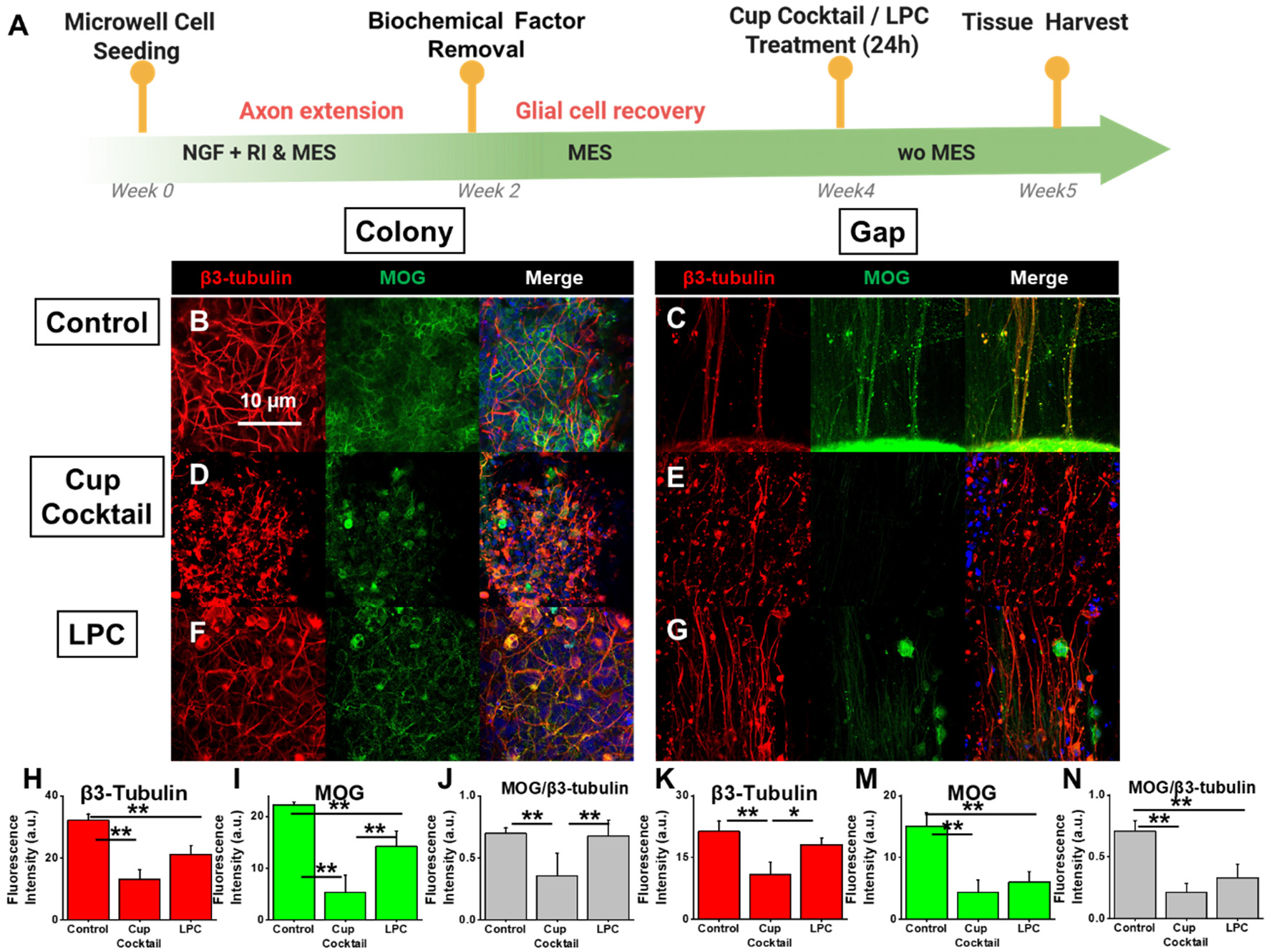
Demyelination of engineered nerve tissues by biochemical reagents. (A) A schematic describing the timeline of engineered nerve tissue formation with myelinated axon extension under mechano-electrical stimulation, biochemical treatment for 24 hrs to induce demyelination, followed by 6 days of normal culture. (B-G) Expression of neural network (β3-tubulin) and myelin sheath (MOG) and cell nuclei (DAPI) without demyelination (B, C), with Cuprizone Cocktail (D, E; cuprizone (1000 µM) + IFN-γ (50 ng/ml) + TNFα (50 ng/ml)), or with LPC (F, G; 30 ug/ml) treatment within the colonies (B, D, F) and inter-colony gaps (C, E, G). (H-N) Quantification of neural network (β3-tubulin in H, K) and myelination (MOG in I, M) and myelination per axon by ratio of MOG/β3-tubulin (J, N) within the colonies (H-J) and inter-colony gaps (K-N) from B-G (n=5, \**p* < 0.05, \*\**p* < 0.01).

### Electrophysiological assessment of demyelinated nerve tissues

To investigate the electrophysiological relevancy of these in vitro demyelination models for neurodegenerative conditions, we conducted MEA measurements on engineered nerve tissues with different conditions of demyelination to characterize neural connectivity and functionality. Following the cell/tissue culture conditions described above, the engineered nerve tissues were carefully removed from their culture chambers and placed upside down onto an MEA surface, ensuring the orthogonal cell alignment of the engineered nerve tissue to the electrode layout so that extended axons in the gap crossed the patterned electrodes. A biphasic pulse (−/+500 mV, 200 µs duration, 10 repetitions) was applied to a row of electrodes, while recording the corresponding responses in the other electrodes to evaluate neural connectivity, transmission distance, and velocity. Representative electrode response maps of extracellular neuronal activity are shown in Fig. 5A, where red circles denote the stimulated electrodes, and gray circles represent the ground electrodes. Other electrode locations display measured action potentials in response to stimulation, reflecting network connectivity. Heatmaps in Fig. 5B illustrate the intensity and distribution of these signals, indicating the locations of responding electrodes and the amplitudes of their responses. The Control group featured a robust neuronal network, with multiple electrodes responding and producing high-intensity signals. In contrast, the Cup Cocktail group exhibited fewer responding electrodes and lower-amplitude signals, primarily localized near the stimulus electrode, suggesting that axonal damage and demyelination severely hinder neural connectivity and reduce signal propagation, corroborating with the IF results (Fig. 4). Although the LPC group also demonstrated diminished responses relative to the Control, its signal response extended farther from the stimulus sites than the Cup Cocktail group, consistent with the IF results in Fig. 4, which showed that the application of LPC preserved the integrity of axons even undergoing demyelination.

**Figure 5.**
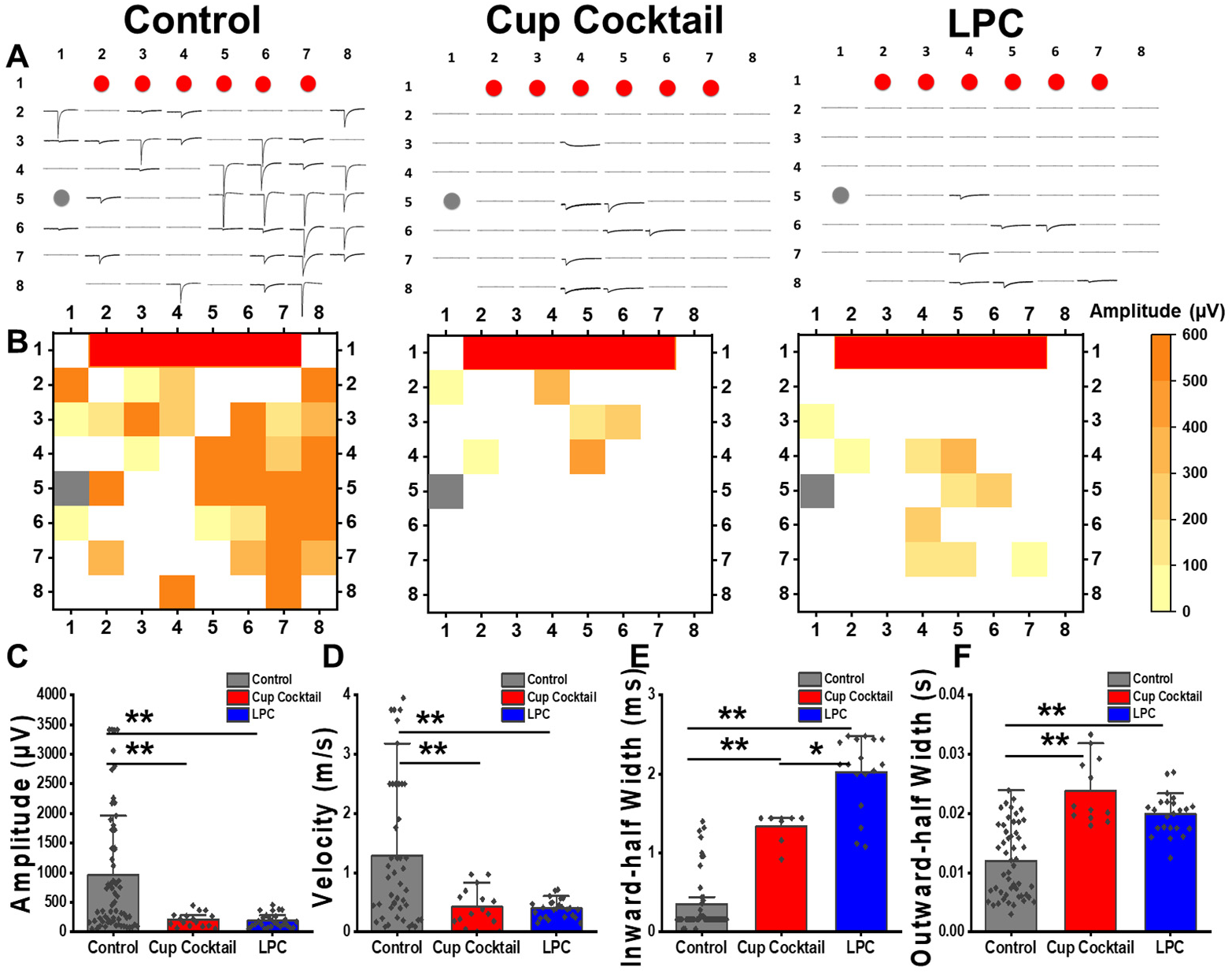
Electrophysiological assessment of engineered nerve tissues with various degrees of myelin damage. (A) Representative electrode maps of extracellular neuronal activities in healthy control and damaged engineered tissues by the application of Cup Cocktail or LPC, determined by a microelectrode array (MEA; 60 electrodes with 200 µm spacing). (B) Action potential heatmaps from (A). (C-F) Quantification of actional potentials, including (C) depolarization amplitude, (D) response velocity, (E) the inward-half width and (F) the outward-half width (n=3; \**p* < 0.05, \*\**p* < 0.01).

Quantitative measurements of action potential amplitude and conduction velocity supported the observation that both Cup Cocktail and LPC demyelination methods compromised neural network functionality (Fig. 5C-F). While the Control group averaged ∼1000 µV in signal amplitude and ∼1.2 m/s in conduction velocity, both of the demyelinated groups displayed significantly compromised amplitudes near 200 µV and velocities ∼ 0.4 m/s (Fig. 5C, D). The demyelination groups both had lower signal amplitude likely due to the impairment in saltatory conduction with the myelin loss [48]. The conduction velocities observed in both the healthy control and demyelination groups were consistent with findings from previous animal studies [49–51]. Additionally, the increased inward- and outward-half-width values in demyelinated samples are likely due to delayed signal initiation and recovery, corresponding to the loss of myelin sheath (Fig. 5E, F) [49, 51, 52]. Overall, the results demonstrate that the Cup Cocktail group exhibited both demyelination and axonal degeneration, whereas the LPC group more selectively induced myelin loss with relatively well-preserved axons in vitro. These two treatments, therefore, provide complementary platforms for modeling the pathology of demyelinating diseases in the SC, enabling targeted investigation of demyelination and axonal loss either independently or in combination.

## 4. Conclusion

The pathology of demyelinating diseases in the SC remains underexplored partly due to the lack of advanced platforms that accurately recapitulate SC anatomy and physiology. In this study, we developed a robust, reproducible in vitro platform that leverages the multiphenotypic differentiation potential of hNSC, microwell technology, and mechano-electrical stimulation from piezoelectric polymeric scaffolds to biofabricate the long, uniaxially aligned, myelinated axons, characteristic of the SC. To model demyelination with or without axonal degeneration, we employed multiple toxin-induced methods, including cuprizone combined with inflammatory cytokines, and LPC. Our platform offers the flexibility to induce distinct pathological features using alternative toxins, demonstrating strong electrophysiological relevance to animal models by demonstrating reduced neural connectivity and impaired signal propagation consistent with axonal injury and demyelination. Overall, this versatile system holds great promise for advancing the study of SC demyelination pathophysiology, disease progression, and the development of potential therapeutic strategies.

## Acknowledgments

L.J. and Y.T. were supported by a TRANSCEND fellowship from the California Institute for Regenerative Medicine (EDUC4–12752). The contents of this publication are solely the responsibility of the authors and do not necessarily represent the official view of CIRM or other agencies of the State of California.

## Data Availability statement

All data analyzed during this study are available from the corresponding author on reasonable request.

## Author contributions

Lu Jin: Writing – review & editing, Writing – original draft, Visualization, Validation, Methodology, Investigation, Formal analysis, Data curation, Conceptualization. Natasha Brinkley: Visualization, Validation, Methodology, Investigation, Formal analysis, Data curation, Conceptualization. Youyi Tai: Methodology, Investigation, Data curation. Galilea Flores: Data curation. Jin Nam: Writing – review & editing, Writing – original draft, Supervision, Resources, Project administration, Investigation, Funding acquisition, Data curation, Conceptualization.

