## supplemental figures for "Engineering an in vitro model of demyelinated spinal cord tissue"

### Supplemental Information

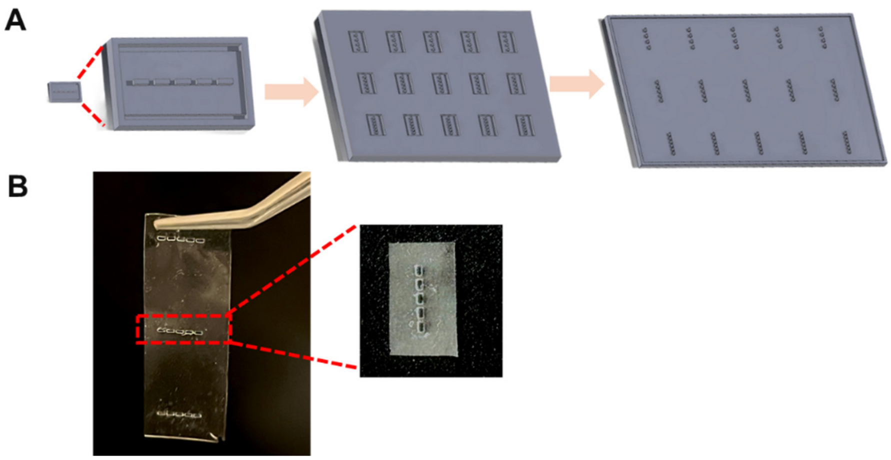


**Figure S1. Design iteration of 3D printed resin molds for the casting of PDMS microwell molds.** (a) Iterative design process of 3D-printed resin molds. The initial prototypes consisted of individually assembled micro-molds bracketed by railed inlays for scaffold fixation. Five modules were created in a single row for repeats, and varying micro-mold distances were created in rows, including 500, 1000, 1500, and 2000 µm. The final design version further simplified bulk fabrication by removing inlays and enabled easier PDMS peeling. (b) Photograph of the final PDMS microwell molds.

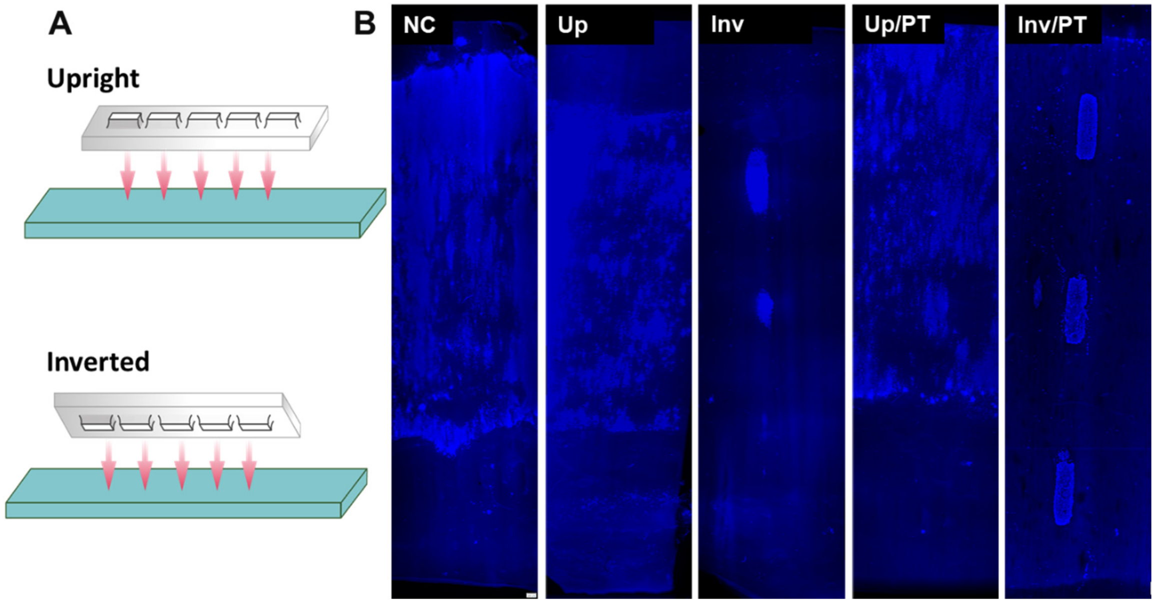


**Figure S2. Optimization of PDMS microwell adhesion to electrospun PVDF-TrFE scaffold.** (A) Illustration of the different sides of the PDMS microwell adhesion to the scaffold by the upright (UP; attaching the side of PDMS microwell mold, which interfaced the 3D printed resin mold during PDMS casting, to the scaffold) or inverted (Inv; attaching the side of PDMS microwell mold, which interfaced air during PDMS casting, to the scaffold) side. The mold side attached to the scaffold was also treated with plasma treatment (PT). (B) Effects of PDMS microwell/scaffold adhesion on cell inoculation, including: 1) NC (negative control, without using the microwell mold, 2) Up (the microwell mold adhered to the scaffold by the upright side without PT), 3) Inv (the microwell mold adhered to the scaffold by the inverted side without PT), 4) Up/PT (the microwell mold adhered to the scaffold by the upright side with PT), 5) Inv/PT (the microwell mold adhered to the scaffold by the inverted side with PT).

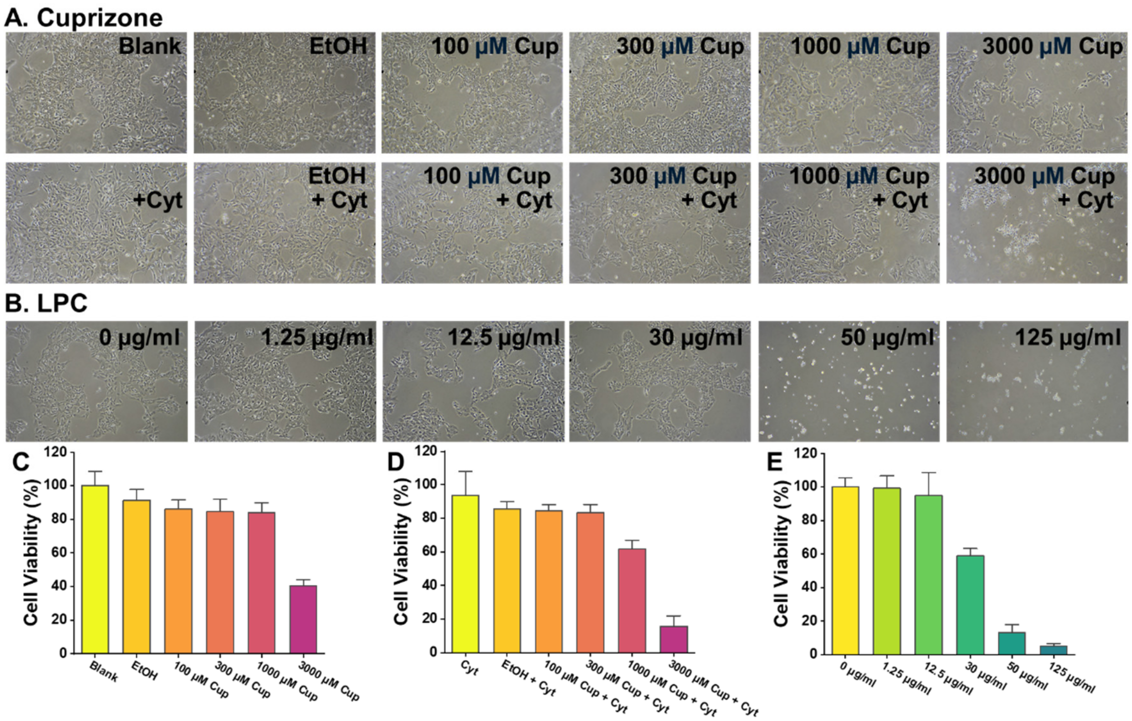


**Figure S3. Optimization of cuprizone and LPC concentration for demyelination.** (A) The cell viability after cells were treated with various concentrations of cuprizone dissolved in 50% ethanol together with or without inflammatory cytokines (IFN-γ 50 ng/ml + TNFα 50 ng/ml) for 24 hrs. The control groups included cells without any treatment, cells with only ethanol (carrier solvent for cuprizone) treatment, and cells with cytokines only. (B) The cell viability after cells were treated with various concentrations of LPC. (C-D) Quantification of cell viability for cuprizone conditions from (A) with (C) or without (D) inflammatory cytokines. (E) Quantification of cell viability for LPC conditions from (B) (n=3).

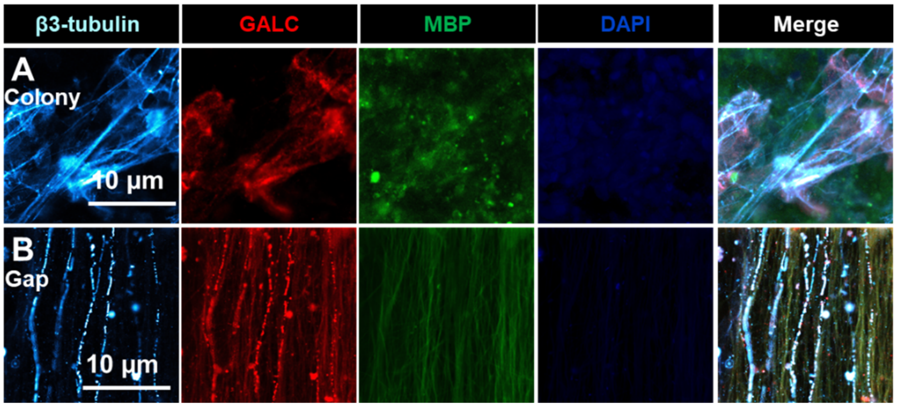


**Figure S4. Effects of cuprizone on engineered nerve tissues.** The application of cuprizone at 3000 μM did not affect nerve structures within colonies (A) and inter-colony gaps (B), assessed by β3-Tubulin, GALC and MBP expression.
